# Mechanistic dissection of BLTP2 targeting to ER-PM contact sites

**DOI:** 10.64898/2026.04.12.718032

**Authors:** Sarah D. Neuman, Amy T. Cavanagh, Nicole Sheffels, Elizabeth Conibear, Arash Bashirullah

## Abstract

The bridge-like lipid transfer proteins (BLTPs) are a novel superfamily of rod-shaped lipid transporters that engage in bulk non-vesicular movement of lipids at organelle membrane contact sites. The molecular and cellular functions of these proteins are still emerging; however, it is clear that one key aspect that regulates BLTP function is targeting to the appropriate membrane contact site(s). Here, we use *Drosophila* as a model system to dissect the mechanisms that drive targeting of *BLTP2* (*hobbit* in *Drosophila*) to endoplasmic reticulum-plasma membrane (ER-PM) contact sites. We demonstrate that a conserved adapter protein, which we name *bilbobaggins* (*bbo*), is required for targeting of Hobbit to ER-PM contacts; importantly, loss of *bbo* phenocopies loss of *hobbit*, indicating that *bbo* is required for *hobbit* function. Additionally, our structure-function analyses show that *cis*-acting sequences in the C-terminal tail of Hobbit are also required for ER-PM targeting. Crucially, our data indicates that that these *cis*-acting sequences and Bbo binding are independent and likely sequential mechanisms that we propose function like a “hook” and “latch” to govern Hobbit targeting. Thus, we define a new regulatory paradigm governing targeting of BLTPs to membrane contact sites.

## INTRODUCTION

The division of cellular compartments into discrete membrane-bound organelles is a defining characteristic of eukaryotic cells. The lipid composition of each organelle membrane is distinct and plays an essential role in regulating organelle identity and function; thus, it is essential that cells carefully control the subcellular distribution of lipids (Van Meer et al., 2008). Lipids can be transported through vesicular or non-vesicular pathways, but the primary mechanism used to move lipids intracellularly is non-vesicular transport. This process is carried out by lipid transfer proteins (LTPs) at membrane contact sites, functional interfaces where two or more organelle membranes come into close (∼10-30 nm) proximity (Prinz et al., 2020; Reinisch et al., 2025). Most known LTPs are small and are shaped like a box with a lid; the box is lined with hydrophobic residues and can accommodate the acyl chains of a single lipid (Wong et al., 2019, 2017). In this manner, these box-like LTPs transfer lipids one at a time from donor to acceptor membranes at organelle membrane contact sites.

The bridge-like lipid transfer proteins (BLTPs) are a novel family of lipid transporters that engage in a different form of lipid transfer. These large (∼2000-4000 aa) proteins are shaped like a tube that is lined with hydrophobic residues (Osawa et al., 2019; Neuman et al., 2022b; Kumar et al., 2018; Li et al., 2020; Valverde et al., 2019; Hanna et al., 2023, 2022; Neuman et al., 2021a; Levine, 2022; Li et al., 2025; Maeda et al., 2019). Recent cryoEM studies (Kang et al., 2025; Hu et al., 2026) and molecular dynamics simulations (Srinivasan et al., 2024) show that BLTPs can harbor tens of lipids at a time. In this manner, the BLTPs act like lipid superhighways to rapidly move lipids from donor to acceptor membranes at organelle membrane contact sites (Neuman et al., 2022b; Hanna et al., 2023; Prinz and Hurley, 2020). There are five basic members of the BLTP superfamily, some of which have multiple paralogs in humans: *BLTP1*, *BLTP2*, *BLTP3A-B*, *ATG2A-B*, and *VPS13A-D* (Braschi et al., 2022). The molecular and cellular functions of these proteins are still emerging, but growing evidence suggests that these proteins are particularly important during processes that require rapid *de novo* synthesis of membrane (Park and Neiman, 2012; Kang et al., 2023), such as autophagosome biogenesis (Maeda et al., 2019; Valverde et al., 2019) or lysosomal damage repair (Hanna et al., 2025; Wang et al., 2025). BLTPs have also been implicated in the regulation of membrane composition during stress responses (Wang et al., 2022; Toulmay et al., 2022; Banerjee et al., 2025; John Peter et al., 2022). Notably, mutations in several of the BLTPs are associated with neurodevelopmental and neurodegenerative diseases in humans (Gueneau et al., 2018; Ueno et al., 2001; Koh et al., 2020; Kolehmainen et al., 2003; Schormair et al., 2018).

Targeting to the appropriate organelle membrane contact site(s) is an essential prerequisite for BLTPs to engage in bulk lipid transfer. The N-terminus of several BLTPs interacts with the endoplasmic reticulum (ER) membrane, either via an N-terminal helical anchor, as seen in *BLTP1* and *BLTP2* (Wang et al., 2022; Neuman et al., 2022a), or by binding of an FFAT motif to ER-resident VAP proteins, as seen in *VPS13A, C*, and *D* (Kumar et al., 2018; Guillén-Samander et al., 2021). The C-terminus of BLTPs, in turn, interacts with other organelle membranes, allowing these proteins to bridge the gap between the ER and another organelle at membrane contact sites. The mechanisms that confer specificity in targeting of BLTPs to membrane contact sites are still emerging, but seem to involve interactions with organelle-specific adapter proteins (Bean et al., 2018; Guillén-Samander et al., 2021; Maeda et al., 2019; John Peter et al., 2017; Cai et al., 2022; Li et al., 2025; Kumar et al., 2018) and/or intrinsic sequences that interact with specific proteins or lipids on organelle membranes (Neuman et al., 2022a; Adlakha et al., 2022; Cai et al., 2022)

We have previously characterized the *Drosophila* ortholog of *BLTP2*, which we named *hobbit*, and we found that *hobbit* mutant animals exhibit a dramatic small body size and developmental arrest during metamorphosis (Neuman and Bashirullah, 2018). *hobbit* function is required for regulated exocytosis in several tissues, including those comprising the endocrine axis that regulates insulin/insulin-like growth factor release (Neuman and Bashirullah, 2018), an essential regulator of systemic growth and body size in all animals (Texada et al., 2020). *hobbit/BLTP2* also has a functional role during synaptogenesis (Neuman et al., 2025; Ito et al., 2026). Work by us and others has found that *hobbit* and its orthologs in yeast localize to ER-plasma membrane (ER-PM) contact sites (Neuman et al., 2022a; Toulmay et al., 2022; Banerjee et al., 2025), and C-terminal truncation of *hobbit* disrupts this localization (Neuman et al., 2022a). Recent studies by us and others also implicate a conserved adapter protein (*HOI1* in *S. cerevisiae; EEIG1* in mammals) in regulating localization of *BLTP2* to ER-PM contact sites (Dziurdzik et al., 2026; Dai et al., 2025).

Here we characterize the function of this novel adapter protein in *Drosophila*. We examine the developmental and cellular functions of *bilbobaggins* (*bbo*), the *Drosophila* ortholog of *HOI1/EEIG1*, and find that they phenocopy those of *hobbit*, indicating that *bbo* is required for *hobbit* function. We also demonstrate that loss of *bbo* disrupts Hobbit targeting to ER-PM contacts. Accordingly, Bbo localizes to the plasma membrane, and overexpression of Bbo is sufficient to increase targeting of Hobbit to ER-PM; this is mediated by binding of Bbo to a conserved α-helical motif in the Hobbit protein (the BLTP2/Hobbit “handle”). In parallel, we examine the role of C-terminal sequences in targeting of Hobbit to ER-PM contacts, and we find that even small truncations disrupt protein targeting and function. However, importantly, overexpression of Bbo is sufficient to recruit C-terminally truncated Hobbit to ER-PM contacts. Together, these results demonstrate that targeting of BLTP2 to ER-PM contact sites is independently regulated by *cis*-acting protein sequences and by binding to *trans-*acting adapter proteins.

## RESULTS

### Loss of *bilbobaggins* (*bbo*) phenocopies loss of *BLTP2/hobbit*

A recent study used a novel affinity purification approach to map nearly the entire *S. cerevisiae* interactome (Michaelis et al., 2023). Interestingly, when Fmp27, one of the yeast orthologs of *BLTP2/hobbit*, was used as bait in this analysis, only one protein (besides Fmp27 itself) was significantly enriched: Ybl086c (Fig. 1A). *YBL086C* was recently named *HOI1* (Dziurdzik et al., 2026). *HOI1* is highly conserved among yeast, flies, and humans, with all three proteins sharing a similar predicted structure consisting of an N-terminal C2 (NT-C2) domain, a disordered “linker” region, and C-terminal α-helices or β-strands (Fig. 1B). Ubiquitous RNAi knockdown of the fly ortholog of *HOI1*, *CG8671*, resulted in a reduction in body size (Dziurdzik et al., 2026) and developmental arrest during metamorphosis (Fig. 1C), highly reminiscent of the phenotypes seen upon ubiquitous RNAi knockdown of *hobbit* (Neuman and Bashirullah, 2018; Dziurdzik et al., 2026) (Fig. 1C). Thus, we name this gene *bilbobaggins* (*bbo*), in honor of the short-statured hobbit character in J.R.R. Tolkien’s novels.

**Figure 1.**
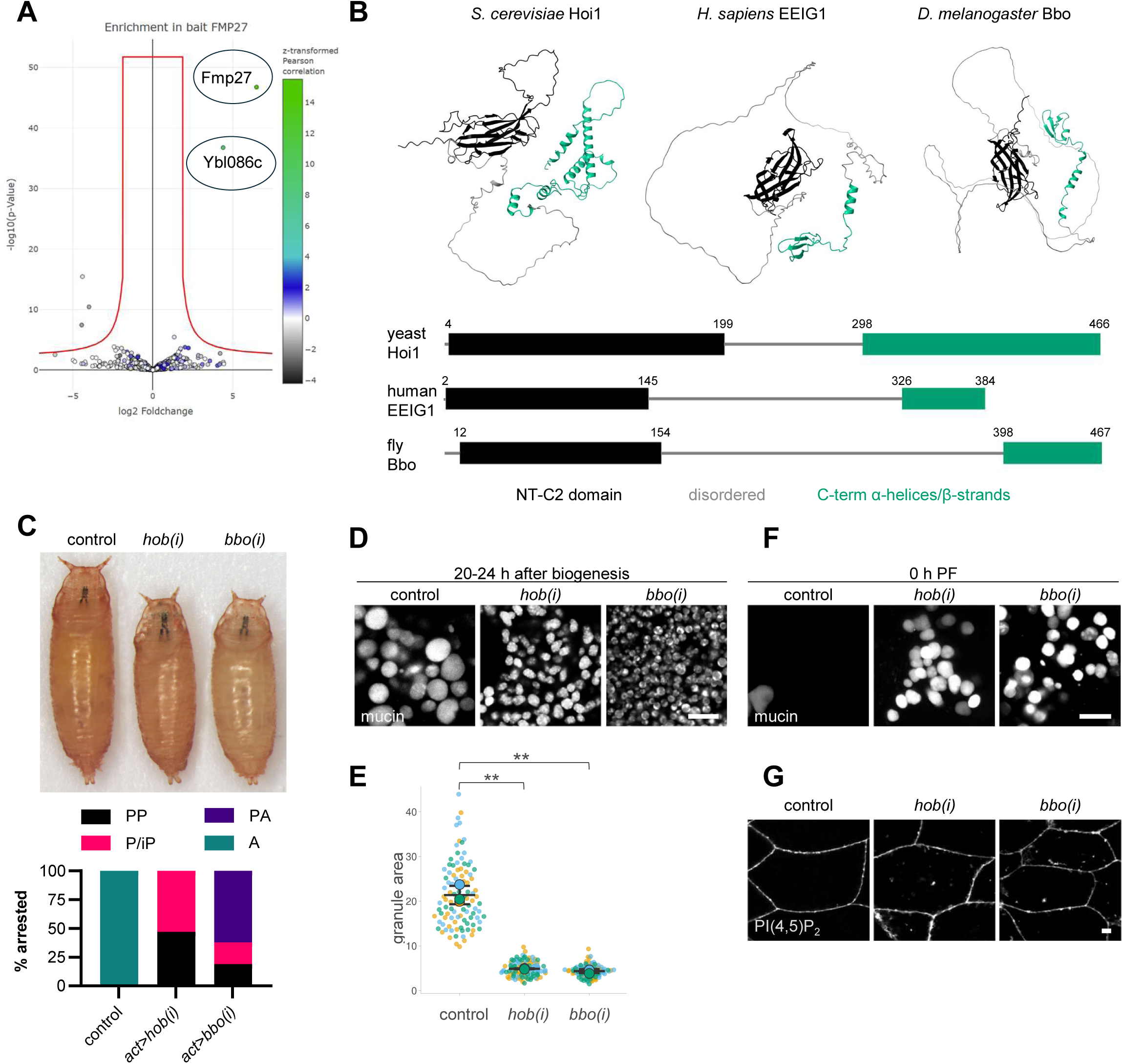
Identification and phenotypic analysis of *bilbobaggins* (*bbo*). **(A)** Volcano plot depicting protein interactors with the yeast ortholog of *hobbit*, Fmp27. Figure modified from www.yeast-interactome.org (Michaelis et al., 2023). **(B)** AlphaFold predicted structures and domain architecture of *S. cerevisiae* Hoi1, *H. sapiens* EEIG1, and *D. melanogaster* Bbo. Boundaries of the NT-C2 domain for each protein were obtained from Uniprot (Bateman et al., 2025). **(C)** Top- representative image of pupa body size in control (*act>luciferase*), ubiquitous *hob* knockdown (*act>hob(i)*), and ubiquitous *bbo* knockdown (*act>bbo(i)*) animals. Bottom- lethal phase analysis shows that ubiquitous knockdown of *hob* or *bbo* results in developmental arrest during metamorphosis. *n*=100 animals per genotype. PP, prepupa; P/iP, pupa/incomplete pupa; PA, pharate adult; A, adult. **(D)** Representative images of mucin-containing secretory granule size in control (*Sgs3-GFP/+; Sgs3>/+*), *hob-RNAi* (*Sgs3-GFP/+; Sgs3>/UAS-hob(i)*), and *bbo-RNAi* (*Sgs3-GFP/+; Sgs3>/UAS-bbo-RNAi*) salivary glands at 20-24 h after the onset of mucin granule biogenesis. Note that mucin granules are small upon salivary gland-specific knockdown of both *hob* and *bbo*. **(E)** ‘SuperPlot’ depicting granule area in control, *hob-RNAi*, and *bbo-RNAi* salivary gland cells shows that mucin granule size is significantly reduced upon knockdown of *hob* or *bbo*. Genotypes are as stated in (D) Bars show mean +/- s.d. for *n*=100 granules per genotype. Granule area was measured in three glands from three separate animals; replicates are color-coded and shown as individual data points. Error bars and statistics calculated using SuperPlotsofData (Goedhart, 2021). ** *p*<0.01. **(F)** Representative images of mucin granules in control, *hob-RNAi*, and *bbo-RNAi* salivary glands at the onset of metamorphosis (0 h PF). Nearly all mucin granules are released in control salivary glands at this timepoint, while a substantial number of mucin granules are retained upon salivary gland-specific knockdown of *hob* or *bbo*. Genotypes are as stated in (D). **(G)** Representative images of the PI(4,5)P_2_ reporter PLCδPH-GFP in control (*Sgs3>PLCδPH-GFP/UAS-luciferase*), *hob-RNAi* (*Sgs3>PLCδPH-GFP/UAS-hob(i)*), and *bbo-RNAi* (*Sgs3>PLCδPH-GFP/UAS-bbo(i)*) salivary glands. PI(4,5)P_2_ is strictly present on the plasma membrane in controls but accumulates intracellularly upon RNAi knockdown of *hob* or *bbo*. All images acquired from live, unfixed tissue and are representative of at least three independent experiments with *n≥*10 tissues from independent animals analyzed. Scale bars: 10 µm.

*hobbit* mutant larval salivary gland cells display several intracellular trafficking phenotypes, including impaired maturation and secretion of mucin-containing secretory granules, defective trafficking of secretory granule membrane proteins (Synaptotagmins and SNARES), enlarged endosomal compartments, and intracellular accumulation of the PM-associated phosphatidylinositol phosphate PI(4,5)P_2_ (Neuman and Bashirullah, 2018; Neuman et al., 2022a). Since ubiquitous knockdown of *bbo* phenocopies ubiquitous knockdown of *hobbit*, we next used the stage- and tissue-specific driver *Sgs3-GAL4* (Cherbas et al., 2003) to test if knockdown of *bbo* also phenocopied knockdown of *hobbit* in the larval salivary glands. We saw that mucin granules in both *hob-RNAi* and *bbo-RNAi* cells were significantly smaller than those in controls at 20-24 h after the onset of mucin biogenesis (Fig. 1D-E), the developmental timepoint when mucin granule maturation is complete (Neuman et al., 2021b). Secretion of mucin granules was also impaired in both genotypes (Fig. 1F). This was also true for the other intracellular trafficking phenotypes: the secretory granule membrane protein Synaptotagmin-1 (Syt1) was absent from mucin granule membranes and instead accumulated in large intracellular “clumps” (Fig. S1A), Rab7-positive endosomes were dramatically enlarged (Fig. S1B), and PI(4,5)P_2_ accumulated intracellularly in both *hob-RNAi* and *bbo-RNAi* cells (Fig. 1G). Together, these results confirm that loss of *bbo* fully phenocopies loss of *hobbit* and suggest that *bbo* is required for *hobbit* function.

### Multiple *bbo* isoforms function in *Drosophila*

To further validate the functional role of *bbo*, we used CRISPR/Cas9 to generate a knockout mutant (see Methods section for details). We found that *bbo^KO^* mutant animals arrested development during larval stages (Fig. 2D). Analysis of mouth hook morphology showed that *bbo^KO^* animals arrested development as a mix of first (L1), second (L2), and early third (eL3) instar larvae (Fig. S3A); no animals crossed the mid-third instar transition to enter the wandering L3 (wL3) stage. In contrast, *hobbit* mutant animals primarily arrest development during metamorphosis (Neuman and Bashirullah, 2018). The earlier developmental arrest of *bbo^KO^* animals compared to *hobbit* mutant animals suggests that *hobbit* and *bbo* may be maternally loaded at different levels, or that *bbo* may perform some essential developmental functions that are independent of those performed by *hobbit*.

**Figure 2.**
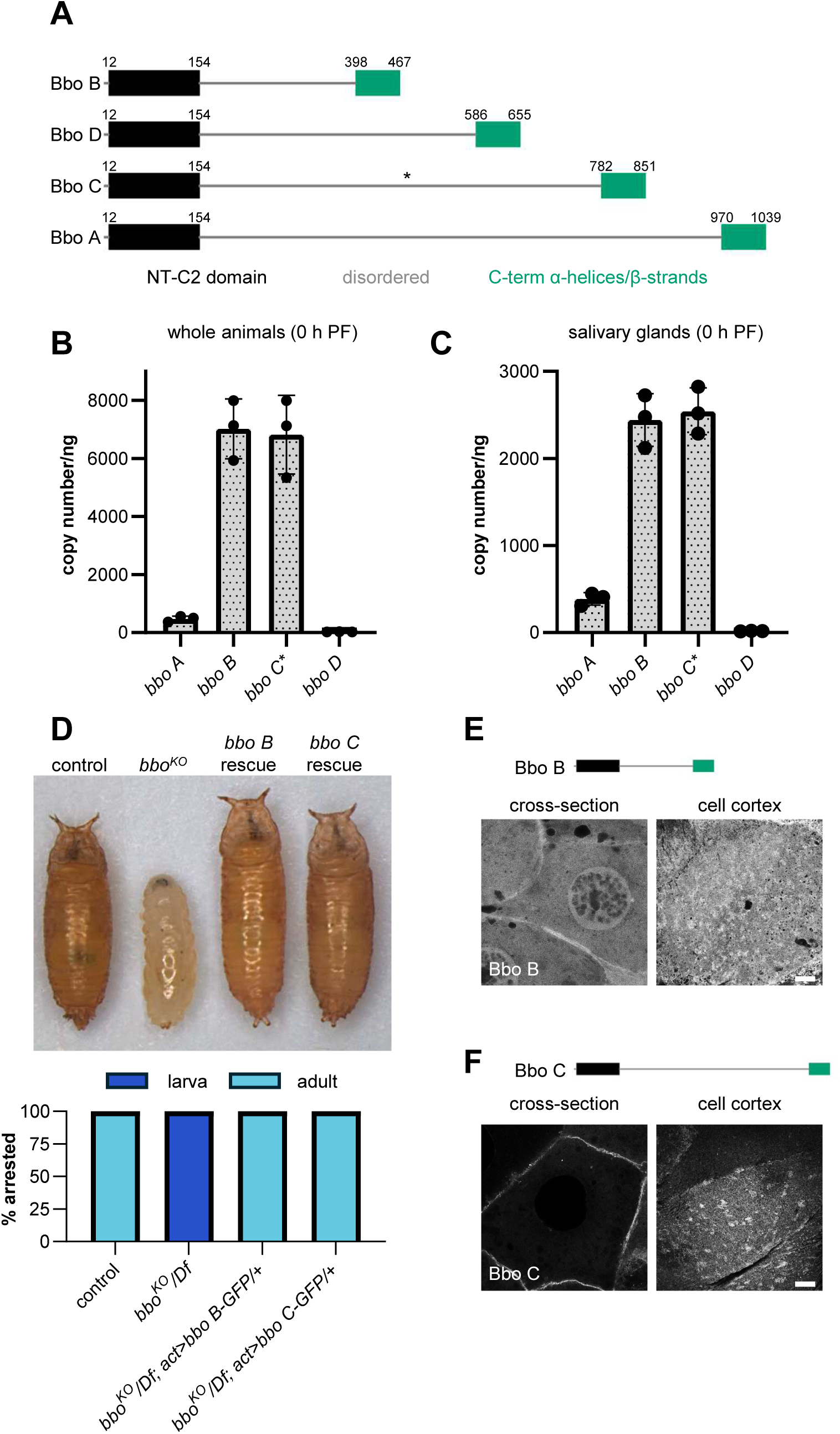
Expression and functional analysis of *bbo* isoforms. **(A)** Schematic depicting the domain architecture and length of predicted *bbo* isoforms in *D. melanogaster*. Boundaries of the NT-C2 domain were obtained from Uniprot (Bateman et al., 2025). Note that the only difference between the isoforms is the length of the disordered “linker” connecting the NT-C2 domain and C-terminal α-helices/β-strands. The asterisk represents slight variability in length of the linker region between the highly similar Bbo isoforms C, E, and F; the linker in isoform E contains 16 additional residues and isoform F contains 26 additional amino acids compared to isoform C (see alignment in Fig. S2). **(B)** Absolute quantification of mRNA transcript levels of *bbo* isoforms A, B, C, and D in whole animals at the onset of metamorphosis (0 h after puparium formation, PF). Note that isoforms C, E, and F cannot be distinguished from one another via qPCR; thus, all three isoforms are measured by the primer set detecting isoform C. Graph shows mean +/- s.d.; individual replicates are overlaid as dots. Samples were collected and analyzed in biological triplicate. **(C)** Absolute quantification of mRNA transcript levels of *bbo* isoforms A, B, C, and D in dissected salivary glands at the onset of metamorphosis (0 h after puparium formation, PF). Note that isoforms C, E, and F cannot be distinguished from one another via qPCR; thus, all three isoforms are measured by the primer set detecting isoform C. Graph shows mean +/- s.d.; individual replicates are overlaid as dots. Samples were collected and analyzed in biological triplicate. **(D)** Top- representative image of body size in control (*w^1118^*), *bbo^KO^* mutant, rescue of *bbo^KO^* mutant with ubiquitous expression of *bbo B*, and rescue of *bbo^KO^* mutant with ubiquitous expression of *bbo C* animals. Bottom- lethal phase analysis shows that *bbo^KO^* mutants arrest development as larvae, while control and rescue animals eclose as viable adults. *n*=50 animals per genotype. **(E)** Representative confocal images of Bbo B-GFP localization in cross-section and at the cell cortex of larval salivary gland cells show that Bbo B localizes throughout the cell but is enriched at the plasma membrane. Images depict a single slice from a z-stack comprising 51 (cross-section) or 41 (cell cortex) slices at a 0.36 μm step size. **(F)** Representative confocal images of Bbo C-GFP localization in cross-section and at the cell cortex of larval salivary gland cells show that Bbo C is strongly enriched at the plasma membrane. Images depict a single slice from a z-stack comprising 54 (cross-section) or 30 (cell cortex) slices at a 0.36 μm step size. All confocal images were acquired from live, unfixed tissue and are representative of at least three independent experiments with *n≥*10 tissues from independent animals analyzed. Scale bars: 10 µm.

*D. melanogaster* is predicted to encode seven mRNA isoforms of *bbo*, labeled A-G, with six distinct proteins produced (isoforms A and G differ only in untranslated regions, not coding sequence; Flybase). The N- and C-termini of all six proteins are the same; they only differ in the length of the disordered “linker” region between the NT-C2 domain and the C-terminal α-helices/β-strands (Fig. 2A; S2). (Note that isoforms C, E, and F differ by only a few amino acids in the linker region; see alignment in Fig. S2A). Bbo isoform B is nearly identical in length and predicted structure to yeast Hoi1 and human EEIG1 (see Fig. 1B). Since exon/intron boundaries are predicted by computational approaches, it is possible that the longer isoforms of *bbo* are not actually expressed *in vivo*. To test this, we designed primers that allowed us to distinguish between isoforms A/G, B, C/E/F, and D (see Fig. S2B) and used qPCR to measure absolute quantities of each transcript, both in whole animals and in dissected salivary glands at the onset of metamorphosis (0 h PF). This analysis showed that isoforms B and C were the most highly expressed, while isoform A was expressed at much lower levels, and isoform D was not expressed at all (Fig. 2B-C). Therefore, at least two isoforms of *bbo*, one short and one long, are abundantly expressed *in vivo*.

We next wondered whether both the short and long Bbo proteins were functional. Thus, we attempted to rescue *bbo^KO^* mutant animals with both isoforms. We generated GFP-tagged overexpression constructs encoding Bbo B or Bbo C, and we found that ubiquitous expression of either protein was sufficient to fully rescue *bbo^KO^*mutants, with all animals eclosing as healthy adults (Fig. 2D). Thus, both Bbo B and Bbo C are functional *in vivo*. Finally, to assess evolutionary conservation of the long Bbo isoforms, we queried HMMER (Potter et al., 2018) using protein sequences specific to isoform A or C (see Fig. S2B), and interestingly, we found that these sequences were only conserved amongst *bbo* orthologs in some species within *Diptera* (Fig. S3B). No homology to other proteins besides *bbo* orthologs was detected. Together, these analyses suggest that Bbo isoform B represents the “ancestral,” functionally complete form of the protein, and the longer isoforms, like isoform C, may have evolved to perform specific functions that are not required in yeast or humans but were evolutionarily selected for in *Diptera*.

### Bbo is required for targeting of BLTP2/Hobbit to ER-PM contacts

As a next step to understanding the function of *bbo*, we used our GFP-tagged overexpression constructs to examine the subcellular localization of the protein in the larval salivary glands. We used live-cell confocal microscopy for these studies. When viewed in cross-section, Bbo B was diffusely present in the cytosol and nucleus but enriched at the plasma membrane of salivary gland cells (Fig. 2E). Bbo C exhibited much stronger enrichment at the plasma membrane, although some signal was still visible in the cytosol (Fig. 2F). When looking specifically at the cell cortex, both Bbo B and Bbo C showed diffuse signal with some locally bright patches, reminiscent of specific subdomains within the plasma membrane, such as ER contact sites (Fig. 2E-F). Therefore, both isoforms of Bbo are enriched at the plasma membrane *in vivo*.

Our phenotypic analysis suggests that *bbo* is required for *hobbit* function. Given that Bbo is enriched at the plasma membrane, and that Hobbit localizes to ER-PM contact sites (Neuman et al., 2022a), we wondered if *bbo* played a role in targeting Hobbit to these sites. Hobbit normally localizes in puncta near the cell cortex of salivary gland cells (Fig. 3A); these puncta strongly co-localize with constitutively active Stim (Stim^DDAA^) (Fig. 3C, E), a marker that we developed to label ER-PM contact sites in flies (Neuman et al., 2022a). Strikingly, we saw a significant reduction in the number of Hobbit cortical puncta upon RNAi depletion of *bbo* (Fig. 3A-B). Accordingly, we also saw a significant reduction in co-localization of Hobbit puncta with Stim^DDAA^ upon knockdown of *bbo* (Fig. 3D-E). These results are consistent with previously reported defects in ER-PM localization of yeast Fmp27 upon loss of *HOI1* (Dziurdzik et al., 2026) and mammalian BLTP2 upon loss of *EEIG1* (Dai et al., 2025).

**Figure 3.**
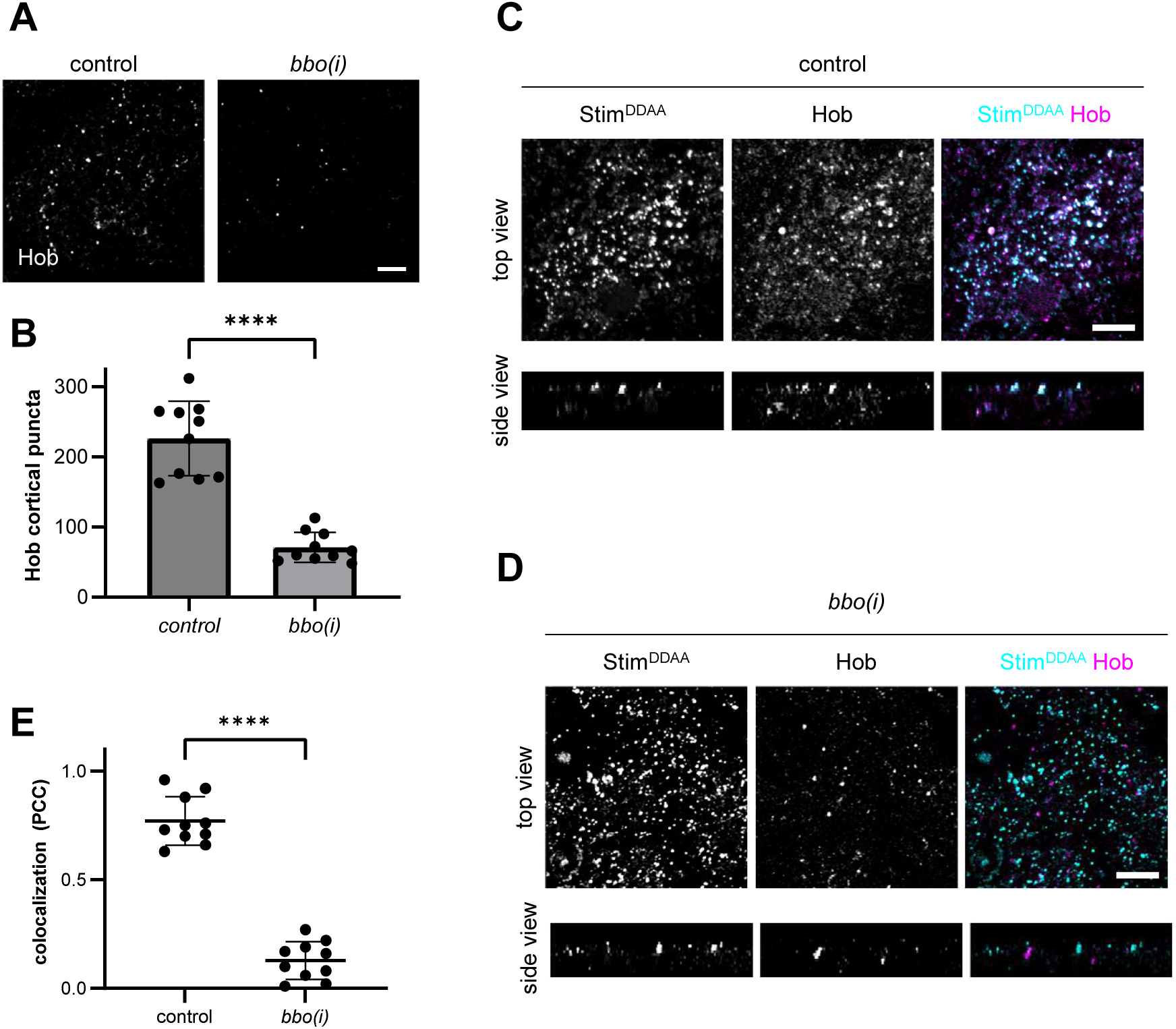
*bilbo* knockdown reduces Hobbit localization to ER-PM contact sites. **(A)** Representative confocal images of cortical Hobbit puncta in control (*Sgs3>hob-mCherry/UAS-luciferase*) and salivary gland-specific *bbo* knockdown (*Sgs3>hob-mCherry/UAS-bbo-RNAi*). Images depict a single slice from a z-stack comprising 20 (control) or 38 (*bbo(i)*) slices at a 0.32 µm step size. **(B)** Quantification of Hobbit cortical puncta in control and salivary gland-specific *bbo* knockdown shows that puncta are significantly reduced in *bbo-RNAi.* Genotypes are as stated in (A). Graph shows mean+/- s.d.; statistics calculated by two-tailed, unpaired *t*-test. *n*=10 salivary glands isolated from independent animals at the onset of metamorphosis (0 h PF); individual data points are overlaid. **(C)** Representative images of the ER-PM marker constitutively-active Stim (Stim^DDAA^-GFP; cyan) and Hobbit-mCherry (magenta) in salivary glands at the onset of metamorphosis (0 h PF). ‘Top view’ images show a maximum intensity projection of 4 slices from a z-stack comprising 24 total slices at a 0.36 µm step size; ‘side view’ images show a transverse (xz) slice through the z-stack. Full genotype: *UAS-Stim^DDAA^-GFP/+; Sgs3>hob-mCherry/UAS-luciferase*. **(D)** Representative images of Stim^DDAA^-GFP (cyan) and Hob-mCherry (magenta) upon salivary gland-specific RNAi knockdown of *bbo*. ‘Top view’ images show a maximum intensity projection of 4 slices from a z-stack comprising 18 total slices at a 0.36 µm step size; ‘side view’ images show a transverse (xz) slice through the z-stack. Full genotype: *UAS-Stim^DDAA^-GFP/+; Sgs3>hob-mCherry/UAS-bbo-RNAi*. All images acquired from live, unfixed tissue and are representative of at least three independent experiments with *n≥*10 salivary glands isolated from independent animals analyzed. Scale bars: 5 µm. **(E)** Quantification of co-localization of Stim^DDAA^-GFP and Hob-mCherry in control and *bbo*-RNAi salivary glands via Pearson’s correlation coefficient shows that co-localization of Stim^DDAA^ and Hobbit is significantly reduced upon knockdown of *bbo*. Genotypes are as stated in (C) and (D). Graph shows mean +/- s.d. from *n=*10 salivary glands isolated from independent animals. Individual data points are overlaid; statistics calculated by two-tailed, unpaired *t*-test. **** *p*<0.0001.

### Green plants do not have a *bbo/HOI1/EEIG1* ortholog

Although *hobbit* is conserved throughout eukaryotes (Fig. 4A; (Neuman and Bashirullah, 2018)), we found that a *bbo* ortholog is completely absent in green plants (Fig. 4A). This intriguing finding led us to examine the predicted structures of Hobbit orthologs in plants to see if there were any obvious differences. Notably, they differed most obviously in a pair of α-helices: in fly Hobbit and human BLTP2, these two helices, which we refer to as the “handle,” are very prominent and project away from the hydrophobic channel (Fig. 4B, S4A), but in *A. thaliana* SABRE, these two helices are intertwined with other helices from another region of the protein (Fig. 4B). The same was true for Hobbit orthologs in *Z. mays*, *O. sativa*, and *P. patens* (Fig. S4A).

**Figure 4.**
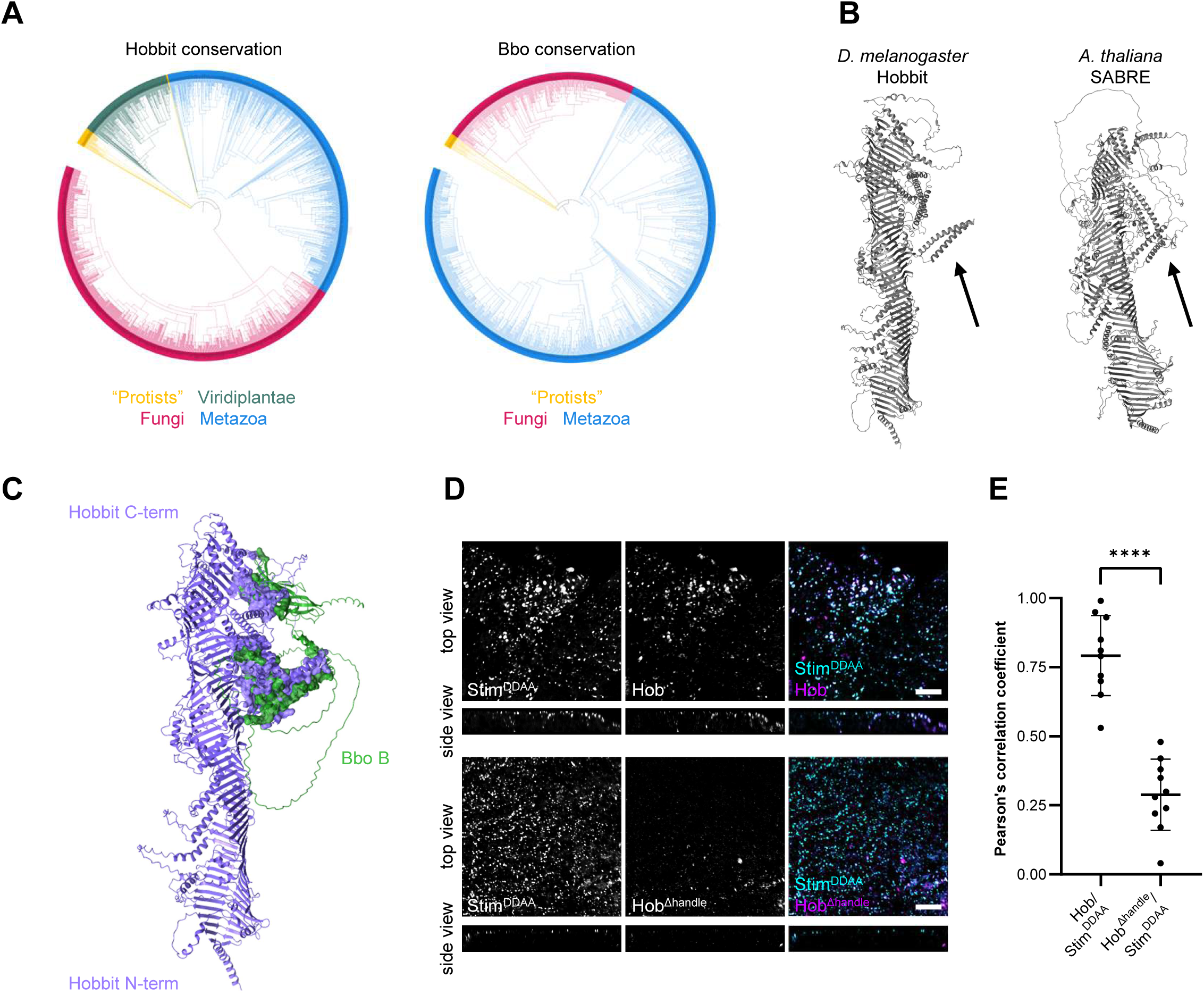
Evolutionary and protein structure analysis of Hobbit and Bbo. **(A)** Phylogenies depicting *hobbit* (left) and *bbo* (right) conservation. Note that *hobbit* is conserved in all major clades within Eukaryota (single-celled eukaryotes (“protists”), plants, fungi, and animals), while *bbo* is missing in plants. **(B)** AlphaFold models of *D. melanogaster* Hobbit and *A. thaliana* SABRE. The black arrow points to the “handle” motif. **(C)** AlphaFold3 modeling of protein-protein interactions between Hobbit and Bbo. Hobbit is shown in purple, and Bbo B is shown in green. Residues that are predicted to physically interact are shown as surface renderings. **(D)** Representative images of Stim^DDAA^-GFP (cyan) and Hob-mCherry (magenta), top, and Stim^DDAA^-GFP (cyan) and Hob^Δhandle^-mCherry, (magenta), bottom in larval salivary gland cells. ‘Top view’ images show a single slice from a z-stack comprising 28 (for wild-type Hobbit) or 24 (for Hobbit^Δhandle^) total slices at a 0.31 µm step size; ‘side view’ images show a transverse (xz) slice through the z-stack. Full genotypes: *UAS-Stim^DDAA^-GFP/+; Sgs3>hob-mCherry/+* and *UAS-Stim^DDAA^-GFP/+; Sgs3>hob^Δhandle^-mCherry/+*. All images acquired from live, unfixed tissue and are representative of at least three independent experiments with *n≥*10 salivary glands isolated from independent animals analyzed. Scale bars: 5 µm. **(E)** Quantification of co-localization of the indicated protein pairs via Pearson’s correlation coefficient shows that wild-type Hobbit-mCherry strongly co-localizes with Stim^DDAA^-GFP, but Hobbit^Δhandle^-mCherry does not. Genotypes are as stated in (D). Graph shows mean +/- s.d. from n=10 salivary glands isolated from independent animals. Individual data points are overlaid; statistics calculated by unpaired, two-tailed *t*-test. **** *p*<0.0001.

We also we used AlphaFold3 (Abramson et al., 2024) to model potential protein-protein interactions between Hobbit and Bbo orthologs. This analysis confidently predicted two interfaces between Hobbit and Bbo: one between the Bbo NT-C2 domain with residues along the hydrophobic groove near the C-terminus of Hobbit, and a second between the C-terminal α-helices/β-strands of Bbo and the “handle” of Hobbit (Fig. 4C). This orientation makes sense, since it would place the Bbo NT-C2 domain (which, for its ortholog EEIG1, has been shown to interact with the PM (Dai et al., 2025)) in close proximity to the PM. In contrast, AlphaFold3-based modeling of *A. thaliana* SABRE with yeast Hoi1, human EEIG1, or fly Bbo did not predict any confident or sensical protein-protein interactions (Fig. S4B). Together, these results point to the α-helical “handle” motif of Hobbit as a potential important binding site for Bbo.

### The conserved Hobbit “handle” is required for Bbo-dependent targeting to ER-PM contact sites

To test a functional requirement for the Hobbit “handle” in Bbo binding, we first compared the localization of wild-type Hobbit to that of Hobbit^Δhandle^, a construct we generated where the handle was seamlessly deleted. The overall predicted structure of Hobbit^Δhandle^ was not substantially different from that of wild-type Hobbit (Fig. 5C). We saw a striking reduction in the number of puncta at the cell cortex with Hobbit^Δhandle^ (Fig. 5C-D), as well as a significant reduction in co-localization with the ER-PM marker Stim^DDAA^ (Fig. 4D-E). Furthermore, ubiquitous expression of *hob^Δhandle^* did not rescue the small body size or metamorphosis lethality of *hobbit* mutant animals (Fig. S4C), indicating that Hobbit^Δhandle^ is not functional. We also examined the co-localization of Bbo with both wild-type Hobbit and Hobbit^Δhandle^. Bbo B and Bbo C both strongly co-localized with Hobbit in puncta at the cell cortex (Fig. 5A, E). Additionally, overexpression of Bbo B or Bbo C significantly increased the number of Hobbit puncta at the cell cortex (Fig. 5A, D), indicating that increased expression of Bbo is sufficient to recruit Hobbit to ER-PM contact sites. In contrast, Hobbit^Δhandle^ did not co-localize with either Bbo B or Bbo C (Fig. 5B, E), and neither Bbo B nor Bbo C were able to recruit Hobbit^Δhandle^ to ER-PM contacts (Fig. 5B, D). Our findings are consistent with other recent reports that point to this motif as a key binding site for Hoi1/EEIG1 (Dai et al., 2025; Dziurdzik et al., 2026). Together, these results demonstrate that the Hobbit “handle” motif is required for *hobbit* function and for Bbo-dependent recruitment to ER-PM contact sites.

**Figure 5.**
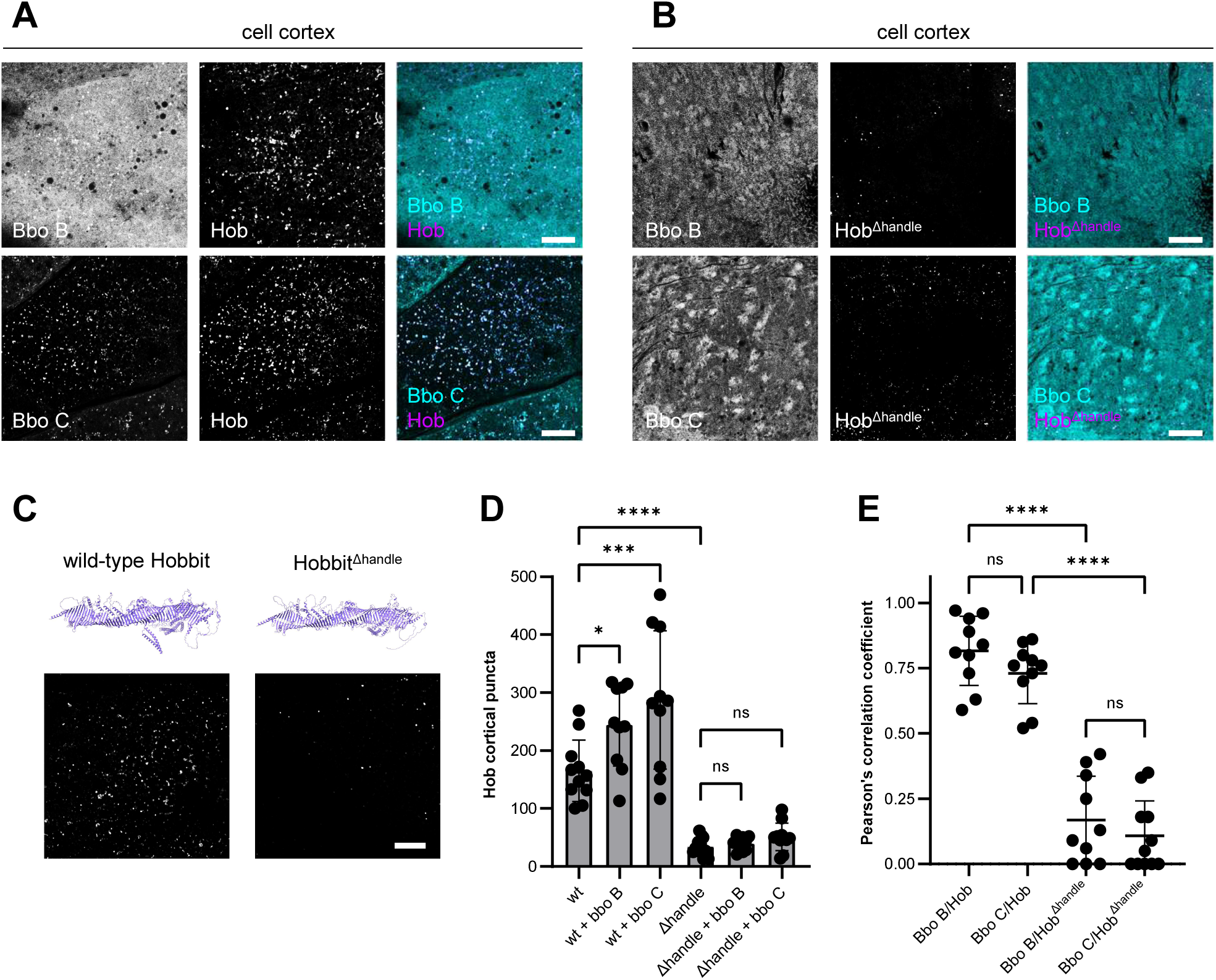
*bbo*-dependent targeting of Hobbit to ER-PM contacts requires the Hobbit “handle.” **(A)** Representative images of Bbo B-GFP (top panels; cyan) or Bbo C-GFP (bottom panels; cyan) with Hobbit-mCherry at the cortex of salivary gland cells at the onset of metamorphosis (0 h PF). Images depict a single slice from a z-stack comprising 23 (top panels) or 21 (bottom panels) slices at a 0.36 µm step size. Full genotypes: *Sgs3>hob-mCherry/UAS-bbo B-GFP* and *Sgs3>hob-mCherry/UAS-bbo C-GFP*. **(B)** Representative images of Bbo B-GFP (top panels; cyan) or Bbo C-GFP (bottom panels; cyan) with Hobbit^Δhandle^-mCherry (magenta) at the cortex of salivary gland cells at the onset of metamorphosis (0 h PF). Images depict a single slice from a z-stack comprising 28 (top panels) or 41 (bottom panels) slices at a 0.36 µm step size. Full genotypes: *Sgs3>bbo B-GFP/UAS-hob^Δhandle^-mCherry* and *Sgs3>bbo C-GFP/UAS-hob^Δhandle^-mCherry*. **(C)** Representative images of Hobbit-mCherry and Hobbit^Δhandle^-mCherry puncta at the cortex of salivary gland cells at the onset of metamorphosis (0 h PF). Images depict a single slice from a z-stack comprising 30 (wild-type) or 31 (Δhandle) slices at a 0.36 µm step size. Full genotypes: *Sgs3>hob-mCherry/UAS-luciferase* and *Sgs3>hob^Δhandle^-mCherry/UAS-luciferase*. AlphaFold predicted protein structures for wild-type Hobbit and Hobbit^Δhandle^ are shown above each confocal image; note that no substantial changes to the overall protein structure are predicted upon deletion of the handle. All images acquired from live, unfixed tissue and are representative of at least three independent experiments with *n≥*10 salivary glands isolated from independent animals analyzed. Scale bars: 10 µm. **(D)** Quantification of Hobbit cortical puncta in the genotypes/conditions shown in (A-C). Note that overexpression of Bbo B or Bbo C increases wild-type, but not Hobbit^Δhandle^, puncta at ER-PM contacts. Hobbit^Δhandle^ shows decreased targeting to ER-PM contacts. Graph shows mean +/- s.d. from n≥10 salivary glands isolated from independent animals. Individual data points are overlaid; statistics calculated by ordinary one-way ANOVA. **(E)** Quantification of co-localization of the indicated protein pairs via Pearson’s correlation coefficient shows that wild-type Hobbit-mCherry strongly co-localizes with Bbo B-GFP and Bbo C-GFP, but Hobbit^Δhandle^-mCherry does not. Genotypes are as stated in (A-C). Graph shows mean +/- s.d. from n≥10 salivary glands isolated from independent animals. Individual data points are overlaid; statistics calculated by ordinary one-way ANOVA. * *p*<0.05, *** *p*<0.001, **** *p*<0.0001. ns = not significant.

### The C-terminal tail of BLTP2/Hobbit is required for targeting to ER-PM contacts

Our previous work demonstrated that deletion of the final 82 amino acids of the Hobbit protein disrupts both ER-PM targeting and protein function (Neuman et al., 2022a). This analysis was informed by one of our *hobbit* mutant alleles, which contains a nonsense mutation just 82 amino acids from the C-terminus of the 2300 amino acid protein (Neuman and Bashirullah, 2018), yet is phenotypically indistinguishable from our other four mutant alleles with nonsense mutations much earlier in the protein (Neuman et al., 2022a). Here, we sought to extend our analysis of the requirement for C-terminal sequences in Hobbit targeting by analyzing the effect of smaller, serial truncations and comparing them to the wild-type protein and Hob^ΔC82^. Notably, the final 82 amino acids of the Hobbit C-terminal tail consist of α-helices and disordered loops that lie entirely outside the hydrophobic groove (Fig. 6C); thus, this “C-terminal tail” is likely highly mobile *in vivo*, making it a prime candidate for a regulatory domain that interacts with the PM to modulate localization and/or function.

**Figure 6.**
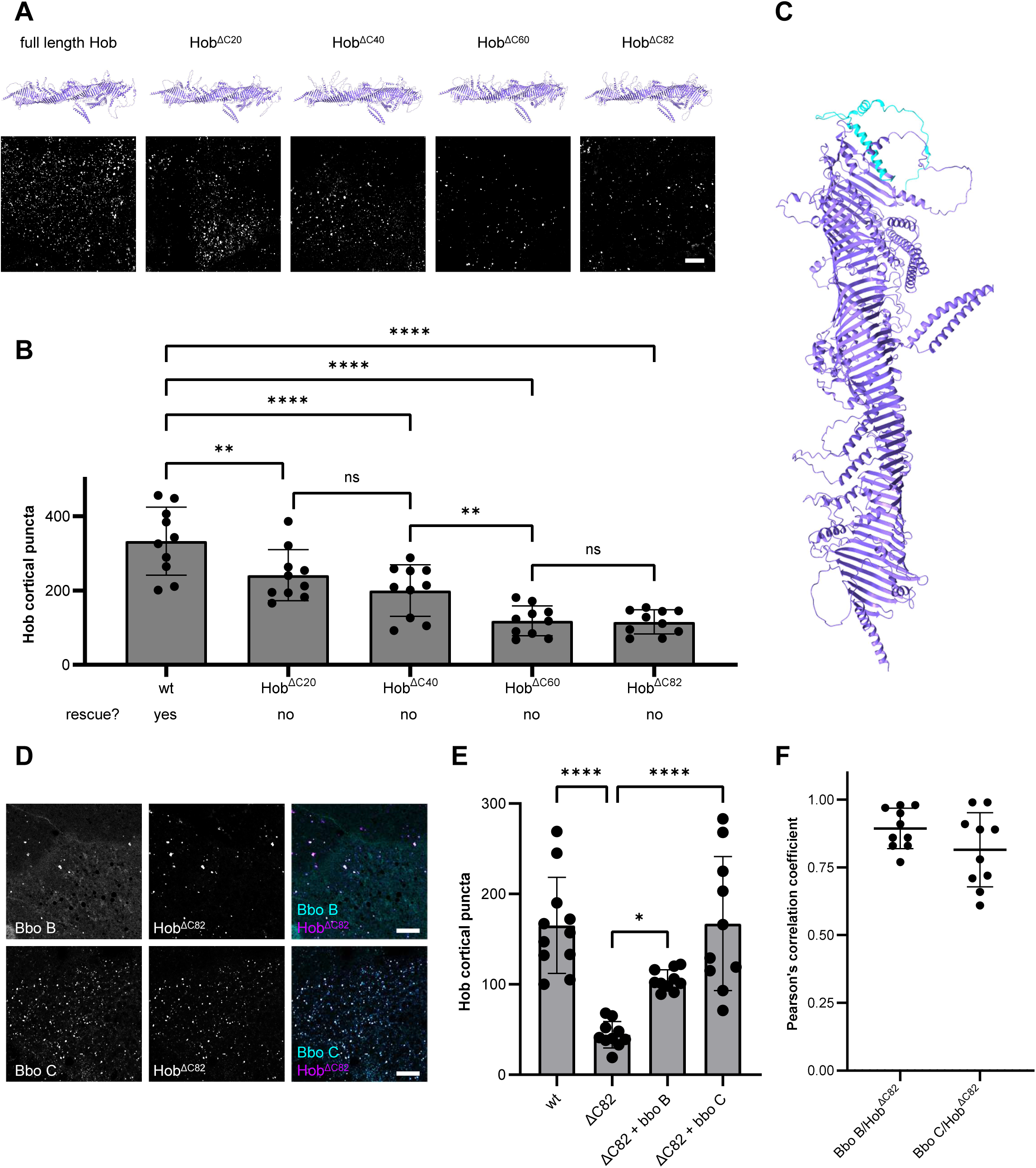
Overexpression of *bbo* can compensate for ER-PM targeting defects upon truncation of the Hobbit C-terminal tail. **(A)** Representative images of wild-type and C-terminally truncated Hobbit-mCherry puncta at the cortex of salivary gland cells at the onset of metamorphosis (0 h PF). Images depict a single slice from a z-stack comprising 18 (wt), 21 (Hob^ΔC20^), 22 (Hob^ΔC40^), 21 (Hob^ΔC60^), or 19 (Hob^ΔC82^) slices at a 0.36 µm step size. Full genotypes: *Sgs3>hob-mCherry/UAS-luciferase*, *Sgs3>hob^ΔC20^-mCherry/UAS-luciferase*, *Sgs3>hob^ΔC40^-mCherry/UAS-luciferase*, *Sgs3>hob^Δ60^-mCherry/UAS-luciferase*, and *Sgs3>hob^ΔC82^-mCherry/UAS-luciferase*. AlphaFold predicted protein structures for wild-type Hobbit and C-terminal truncations are shown above each confocal image. **(B)** Quantification of wild-type and C-terminally Hobbit puncta in the genotypes shown in (A). All C-terminal truncations significantly impair Hobbit targeting to ER-PM contact sites. Bottom row indicates whether each protein can rescue *hobbit* mutant animals (also see Fig. S6). Graph shows mean +/- s.d. from n=10 salivary glands isolated from independent animals. Individual data points are overlaid; statistics calculated by ordinary one-way ANOVA. **(C)** AlphaFold predicted structure of Hobbit; the C-terminal 82 amino acids are colored in cyan. Note that all final 82 amino acids lie outside of the hydrophobic groove. **(D)** Representative images of Bbo B-GFP (top panels; cyan) or Bbo C-GFP (bottom panels; cyan) with Hobbit^ΔC82^-mCherry (magenta) at the cortex of salivary gland cells at the onset of metamorphosis (0 h PF). Images depict a single slice from z-stacks comprising 27 slices at a 0.36 µm step size for both the top and bottom panels. Full genotypes: *Sgs3>bbo B-GFP/UAS-hob^ΔC82^-mCherry* and *Sgs3>bbo C-GFP/UAS-hob^ΔC82^-mCherry*. All images acquired from live, unfixed tissue and are representative of at least three independent experiments with *n≥*10 salivary glands isolated from independent animals analyzed. Scale bars: 10 µm. **(E)** Quantification of wild-type Hobbit-mCherry and Hob^ΔC82^-mCherry puncta at the cell cortex (images not shown) and Hobbit cortical puncta in the genotypes/conditions shown in (D). Hobbit^ΔC82^ shows decreased targeting to ER-PM contacts; however, overexpression of Bbo B or Bbo C increases Hobbit^ΔC82^ targeting to ER-PM. Graph shows mean +/- s.d. from n≥10 salivary glands isolated from independent animals. Individual data points are overlaid; statistics calculated by ordinary one-way ANOVA. **(F)** Quantification of co-localization of the indicated protein pairs via Pearson’s correlation coefficient shows that Bbo B-GFP and Bbo C-GFP strongly co-localize with Hobbit^ΔC82^-mCherry. Genotypes are as stated in (D). Graph shows mean +/- s.d. from n≥10 salivary glands isolated from independent animals. Individual data points are overlaid. * *p*<0.05, ** *p*<0.01, **** *p*<0.0001. ns = not significant.

We generated a series of new mCherry-tagged constructs that each removed 20 amino acid residues from the Hobbit C-terminal tail: Hob^ΔC20^, Hob^ΔC40^, and Hob^ΔC60^ (Fig. 6A), and tested their ability to target to ER-PM contacts. All the C-terminal truncations significantly reduced the number of puncta at the cell cortex (Fig. 6A-B), indicating that none were properly targeted to ER-PM. Additionally, we found that ubiquitous expression of the truncation constructs failed to rescue *hobbit* mutant animals (Fig. S5), highlighting the critical functional importance of the C-terminal tail. Interestingly, six of the final 20 amino acids of Hobbit have positively charged sidechains (K or R); thus, this region of the protein could act like a “basic patch” (Giordano et al., 2013) that interacts with phospholipid headgroups on the plasma membrane to anchor the protein. This model is consistent with our data, where deletion of the final 20 amino acids of Hobbit is sufficient to disrupt ER-PM targeting, but larger deletions further exacerbate the defect, likely by disrupting other elements that interact with specific lipids or proteins on the plasma membrane.

### *cis-*acting C-terminal tail and *trans*-acting adapter protein binding are independent mechanisms that target BLTP2/Hobbit to ER-PM contact sites

Our data demonstrates that loss of *bbo* or truncation of the Hobbit C-terminal tail independently impair targeting of the protein to ER-PM contacts, raising the question of how, or if, these two mechanisms interact. To begin to answer this question, we tested whether overexpression of Bbo could recruit Hob^ΔC82^ to ER-PM. The “handle” is left intact in Hob^ΔC82^ (Fig. 6A), and Bbo is not predicted to interact with the C-terminal tail (see Fig. 4C); thus, the primary binding site for Bbo is still present in this version of the protein. Strikingly, we found that overexpression of Bbo B or Bbo C significantly increased the number of Hob^ΔC82^ puncta at ER-PM (Fig. 6D-E), nearly restoring puncta counts to wild-type levels. Furthermore, Hob^ΔC82^ strongly colocalized with Bbo B and C (Fig. 6F), indicating that overexpression of Bbo can recruit Hob^ΔC82^ to ER-PM contacts. This data shows that adapter protein (Bbo) binding and the Hobbit C-terminal tail are independent mechanisms that govern targeting to ER-PM contact sites. Additionally, since overexpression of Bbo can compensate for deletion of the C-terminal tail, our data suggests that these two factors likely act sequentially, with adapter protein binding first bringing Hobbit close to the PM, and the C-terminal tail subsequently anchoring Hobbit at ER-PM contact sites.

## DISCUSSION

The BLTPs are mysterious proteins with direct relevance to human health and disease. How these proteins are targeted to their site of action at membrane contact sites remains a fundamental question in the field. Our data presented here supports a model in which BLTP2/Hobbit targeting to ER-PM contacts is governed by two separable but likely sequential inputs: a *trans*-acting adapter protein-dependent element that acts like a “hook” to bring the protein in proximity to the PM, and a *cis*-acting C-terminal tail element that acts like a “latch” to stably anchor the protein at ER-PM contacts. Together, these findings highlight a novel regulatory paradigm governing targeting of BLTPs to membrane contact sites.

In our “hook and latch” model for BLTP2 targeting, Bbo/Hoi1/EEIG1 localizes to the plasma membrane via its NT-C2 domain, and the disordered, flexible “linker” region allows the C-terminal α-helices/β-strands to bind the BLTP2/Hobbit handle, acting as a “hook” that brings BLTP2, which is already anchored in the ER membrane via an N-terminal helix (Neuman et al., 2022a), in close proximity to the plasma membrane. Then, the *cis*-acting C-terminal tail of BLTP2, including a putative basic patch, interacts with lipids on the plasma membrane to act as a “latch,” further stabilizing BLTP2 at ER-PM contact sites. A recent study demonstrated that phosphatidylinositol-4-phosphate (PI4P) is involved in targeting BLTP2 to ER-PM contact sites in mammalian cells (Dai et al., 2025); since this lipid is abundant in the plasma membrane (Balla, 2013), it is plausible that the C-terminal tail primarily interacts with PI4P to stabilize the protein at ER-PM contacts. Additionally, modeling of BLTP2 orthologs consistently places a helix within the C-terminal tail in a position that blocks the opening of the hydrophobic groove; thus, it is possible that “latching” to the PM also triggers a conformational change in the C-terminal tail that makes BLTP2 competent for lipid transfer. Whether this model of sequential, bipartite regulation of targeting to a specific membrane contact site is generalizable to other BLTPs is an open question.

Our comparison of the predicted structures of *BLTP2* orthologs revealed intriguing differences that suggest a potential divergence in cellular function between plants and opisthokonts. The overall shape and length of the hydrophobic groove is largely unchanged among *BLTP2* orthologs in plants, fungi, and animals, which strongly suggests that all engage in bulk lipid transfer as their primary molecular function. Accordingly, the reported phenotypes upon mutation of *BLTP2* orthologs in plants center around pollen tube growth, root hair growth, and cell plate deposition (Pietra et al., 2015; Procissi et al., 2003; Xu and Dooner, 2006; Benfey et al., 1993; Aeschbacher et al., 1995; Cheng and Bezanilla, 2021; Pietra et al., 2013), all processes that rely heavily on membrane trafficking pathways to drive rapid expansion or *de novo* synthesis of membrane compartments (Hepler et al., 2001; Boruc and Van Damme, 2015). However, the predicted structure of “adornments,” like the BLTP2 handle, that lie outside the hydrophobic groove look very different between plants and opisthokonts. The reported localization of *BLTP2* is quite consistently to ER-PM contact sites (with few exceptions) in yeast, fly, and mammalian cells (Neuman et al., 2022a; Toulmay et al., 2022; Banerjee et al., 2025; Dziurdzik et al., 2026; Dai et al., 2025; Parolek and Burd, 2024). In contrast, the localization of *BLTP2* orthologs in plants is much less clear, with reported enrichment at the Golgi (Xu and Dooner, 2006), various endomembranes (Pietra et al., 2013), and ER puncta (Cheng and Bezanilla, 2021). These findings, coupled with the absence of a *bbo* ortholog in plants, suggest that plants likely evolved different regulatory mechanisms that govern their subcellular localization and thus their cellular function.

Although Bbo isoform B is broadly conserved among opisthokonts, isoform C is restricted to select *Dipteran* lineages. Homology searches using isoform C–specific sequences recovered only Bbo orthologs and no other related proteins, indicating that these isoforms with long linkers are unique to *Diptera*. This raises the question of why they emerged and were retained in flies and mosquitoes. One possibility relates to *Diptera*-specific biology, particularly rapid larval growth. For example, *D. melanogaster* increases body mass by ∼200 fold over just four days of larval development (Tennessen et al., 2011). Our data suggests that Bbo C may enhance recruitment of Hobbit to ER–PM contact sites, potentially supporting elevated lipid transfer to meet the demands of rapid cellular and organelle expansion. Alternatively, Bbo C may have Hobbit-independent functions that confer fitness advantages during development or reproduction. Resolving these possibilities will require the generation and analysis of isoform-specific *bbo* mutants.

Other BLTPs certainly contain “adornments” akin to the Hobbit handle on the hydrophobic groove; indeed, VPS13A contains four such elements that all lie near its C-terminus: the VPS13 adapter binding (VAB) domain, a pair of helices called the ATG_C domain, a pleckstrin homology (PH) domain, and a WWE domain (Levine, 2022). Organelle-specific adapter proteins that bind to the VAB domain have been identified (Bean et al., 2018; Park et al., 2013, 2021), and a recent cryoEM study shows that the VPS13A PH domain binds to the plasma membrane-localized scramblase XKR1 (Hu et al., 2026). This same study also suggests that the VAB domain can adopt different conformations that either block or open the hydrophobic groove, thereby regulating the competence of the protein for lipid transfer (Hu et al., 2026). Thus, multifactorial control of localization and function appears to be a common feature of BLTPs, though future studies will be required to determine if these mechanisms act independently and/or sequentially like we observe for BLTP2. Notably, independent targeting mechanisms also provide opportunities for layers of regulation that could up or downregulate targeting and/or activity of BLTPs during specific developmental windows or in response to specific stimuli. Additional work will be required to parse out the larger biological relevance and impact of targeting mechanisms for BLTPs.

## ACKNOWLEDGEMENTS

Stocks obtained from the Bloomington Drosophila Stock Center (NIH P40OD018537) and plasmids obtained from the Drosophila Genomics Resource Center (NIH 2P40OD010949) were used in this study. This work was supported by funding from the National Institutes of Health (1R01GM155154 to A.B.), the Canada Foundation for Innovation (Leading Edge Fund 30636), and Canadian Institutes of Health Research (OGB-177941 and PJT-180544 to E.C.).

## METHODS

### *Drosophila* stocks and husbandry

The following stocks were obtained from the Bloomington *Drosophila* Stock Center: *w^1118^* (RRID:BDSC_3605), *Sgs3-GFP* (RRID:BDSC_5884), *Sgs3-GAL4* (RRID:BDSC_6870), *hs-GAL4* (RRID:BDSC_1799), *UAS-bbo-RNAi* (RRID:BDSC_41965), *UAS-Syt1-GFP* (RRID: BDSC_6926), *Rab7-EYFP* (RRID:BDSC_62545), *UAS-PLCδPH-GFP* (RRID:BDSC_39693), *UAS-luciferase* (RRID:BDSC_35788), *act-GAL4* (RRID:BDSC_3954), *Df(2L)Exel6048* (RRID: BDSC_7530). The following fly stocks were previously described: *hob^2^* (Neuman and Bashirullah, 2018), *hob^3^* (Neuman and Bashirullah, 2018), *UAS-hob-RNAi* (Neuman and Bashirullah, 2018), *UAS-hob-mCherry* (Neuman et al., 2022a), *UAS-hobΔC82-mCherry* (Neuman et al., 2022a), *UAS-Stim^DDAA^-GFP* (Neuman et al., 2022a). *Sgs3-DsRED* was kindly provided by A. Andres (University of Nevada, Las Vegas, NV, USA) (Costantino et al., 2008). The RNAi knockdown efficiency of *UAS-hobbit-RNAi* and *UAS-bbo-RNAi* was previously reported (Neuman and Bashirullah, 2018; Dziurdzik et al., 2026). All crosses were grown in uncrowded bottles or vials containing standard cornmeal-molasses media (Archon Scientific) and kept in an environmental chamber set to 25°C.

### Pupa imaging, body size quantification, and lethal phase analysis

Pupa images were captured on an Olympus SXZ16 stereomicroscope coupled to either an Olympus DP72 digital camera with DP2-BSW software or an Olympus DP75 digital camera with Olympus cellSens Standard 4.3 software. Pupa pictured together were initially captured in the same image, but animals were individually aligned and rotated post-acquisition to improve image aesthetics. Body size was quantified by measuring the length and width of each pupa in Adobe Photoshop CS6, and pupa volume was calculated using a published formula (Delanoue et al., 2016) For lethal phase analysis, pupae were allowed to age on grape agar plates at 25°C for 1 week; then, Bainbridge and Bownes staging criteria (Bainbridge and Bownes, 1981) were used to determine lethal phase. To image larval mouth hooks, developmentally arrested *bbo^KO^/Df* larvae were firmly squashed between a slide and coverslip, then imaged on a Zeiss Axioskip 2plus coupled to an Olympus DP75 digital camera with Olympus cellSens Standard 4.3 software. Reference photos in (Graf et al., 1992) were used to determine developmental staging.

### Generation of *Drosophila* transgenic and CRISPR/Cas9 mutant lines

To generate the *hobbit* C-terminal deletion constructs, the Gateway cloning entry vector *pENTR1A*-*hobbit* (Neuman and Bashirullah, 2018) was digested with SpeI-HF (New England Biolabs R3133) and NotI (New England Biolabs R0189), and the resulting 290 bp fragment was cloned into *pGEM T-easy* (Promega A1360). The deletions were made by divergent PCR using *pGEM T-easy-hob(SpeI/NotI)* as a template. The primers were as follows: for *hob^ΔC20^* 5’ GTT TTC ATT CAA TAG GCG ATT TCC GAA CAA C 3’ and 5’ TTG CGG CCG CAA TCA CTA GTG AAT TC 3’; for *hob^ΔC40^* 5’ GGG TCG CTC GTT CGC CTG CGG AT 3’ and 5’ TTG CGG CCG CAA TCA CTA GTG AAT TC 3’; for *hob^Δ60^* 5’ GAT GGC CTG CGA GAA TAT CAC CCG CC 3’ and 5’ TTG CGG CCG CAA TCA CTA GTG AAT TC 3’. The resulting PCR products were digested with DpnI (New England Biolabs R0176), phosphorylated with T4 Polynucleotide Kinase (New England Biolabs M0201), and ligated back into *pGEM T-easy* with T4 DNA Ligase (New England Biolabs M0202). Subsequently, *pGEM T-easy hob^ΔC20^*, *pGEM T-easy hob^ΔC40^*, and *pGEM T-easy hob^ΔC60^* were digested with SpeI-HF and NotI and ligated back into *pENTR1A*-*hobbit* (also digested with SpeI-HF and NotI). Each truncation construct was sequence-verified, then recombined into the Gateway destination vector *pBID-UASC-G-mCherry* (Neuman et al., 2022a; Wang et al., 2012) using LR Clonase I (Invitrogen 11791019) to generate the final plasmids *UAS-hob^ΔC20^-mCherry*, *UAS-hob^ΔC40^-mCherry*, and *UAS-hob^ΔC60^-mCherry*. The Hob^Δhandle^ construct removes amino acids 928-1003 of the Hobbit protein. To generate *UAS-hob^Δhandle^-mCherry*, *pENTR1A-hobbit* was used as a template for two PCR reactions: the first to generate a 191 bp fragment immediately upstream of amino acid 920 (primers: 5’ ACG TCC GCG GCA ATT GGA TCC TTT AAA GCC ATT TTC CCG 3’ and 5’ CAG GCC AGC AAT CGG GTG TCG CTA ATT TCC AGC AAG 3’) and the second to generate a 161 bp fragment immediately downstream of amino acid 1003 (primers: 5’ CTT GCT GGA AAT TAG CGA CAC CCG ATT GCT GGC CTG 3’ and 5’ CCG CGG CAC CAA AGT GTA GAG AAC TCG AGA CCC 3’). These two fragments were then used in an overlapping PCR reaction to generate a 352 bp fragment, which was subsequently digested with MfeI-HF (New England Biolabs R3589) and SacII (New England Biolabs R0157) and ligated into *pENTR1A-hobbit*. The resulting plasmid (*pENTR1A-hob^Δhandle^*) was sequence-verified and recombined into the Gateway cloning destination vector *pBID-UASC-G-mCherry* using LR Clonase. To generate *UAS-bbo B-GFP* and *UAS-bbo C-GFP*, the *bbo* isoform B and isoform C coding sequences were obtained from Flybase (Öztürk-Çolak et al., 2024) and were synthesized and cloned via seamless cloning into *pUAS-C-EGFP-BD-attB* (RRID:DGRC_1503) at the XhoI site (GenScript). All final plasmids were sequence-verified and injected into *VK00027* flies for phiC31-mediated site-directed integration using standard techniques (BestGene, Inc.). The *bbo^KO^*mutant was generated by CRISPR-mediated mutagenesis, performed by WellGenetics Inc. using methods modified from (Kondo and Ueda, 2013). In brief, the upstream sequences AAG GCC ATG CCT GGG GCG CT[AGG]/CTT CTT GGG CCT AGC GCC CC[AGG] and the downstream sequences ATT TGT GGA ATG TGG TCG CA[CGG]/ATG CTG CCT ATG AGC TAA TG[AGG] were cloned into U6 promoter plasmids separately. The cassette 3xP3-RFP, which contains a floxed 3xP3-RFP, and two homology arms were cloned into pUC57-Kan as a donor template for repair. *bbo*-targeting gRNAs and hs-Cas9 were supplied in DNA plasmids, together with donor plasmid for microinjection into embryos of the control strain *[LWG132] w[1118]; P{nos-Cas9, y+, v+}3A/TM6B, Tb[1]*. F1 flies carrying the selection marker 3xP3-RFP were further validated by genomic PCR and sequencing. The final *bbo^KO^* allele deletes the coding sequence of *bbo* (-3 nt to +4,835 nt from ATG of *bbo*; a 4,838 bp deletion) and replaces it with the 3xP3-RFP cassette.

### Confocal microscopy and image quantification

All confocal images were acquired using an Olympus FV3000 laser scanning confocal microscope (100x oil immersion objective, NA 1.49) with FV31S-SW software at room temperature. Only live, unfixed tissues were imaged; salivary glands were dissected from animals of the appropriate developmental stage and genotype in PBS, then mounted in 1% low-melt agarose (Apex Chemicals 20-103) made in PBS. Tissues were imaged immediately and for no longer than 15 minutes after mounting. At least 10 salivary glands dissected from independent animals were imaged for each genotype per experiment. Images that were obtained as *z*-stacks, as indicated in the figure legends, were deconvolved using three iterations of the Olympus CellSens Deconvolution for Laser Scanning Confocal Advanced Maximum Likelihood algorithm. Co-localization was quantified by Pearson’s correlation coefficients calculated using Olympus CellSens software. All images showing Hobbit cortical puncta were acquired as *z*-stacks; acquisition began outside of the cells/tissue and finished ∼8 μm inside of the cells/tissue (∼10 μm total depth). For quantification of Hobbit cortical puncta, transverse sections of each image were viewed to identify the single slice at the surface of the cell; then, total Hobbit puncta in that slice were quantified using the Count and Measure function of Olympus CellSens software. Identical thresholding parameters were used for puncta quantification of all images. Mucin granule area was quantified using ImageJ (Schneider et al., 2012); a minimum of 30 granules per image were quantified, and three images obtained from independent animals were used. Mucin granule area was plotted using SuperPlotsOfData (Goedhart, 2021; Lord et al., 2020). For figure display purposes, brightness and contrast of all images were optimized post-acquisition using Olympus FV31S-SW software.

### Phylogenetic analysis

To identify orthologs of *D. melanogaster hobbit* and *bbo* isoform B, the protein sequences were obtained from Flybase (Öztürk-Çolak et al., 2024) and queried against the Uniprot database (2025_01) using HMMER (Potter et al., 2018). Significant hits were then mapped to Uniprot IDs and matched to NCBI Taxonomy IDs (TaxIDs) using the ID mapping function within Uniprot (Bateman et al., 2025). The NCBI TaxIDs were then submitted to phyloT (phylot.biobyte.de) to generate a phylogenetic tree, which was visualized, annotated, and edited using Interactive Tree of Life (Letunic and Bork, 2024). A similar process was followed to identify orthologs of *bbo* isoform A and isoform C, except that the sequences used to query HMMER were specific to each isoform.

### Protein structure prediction and analysis

When available, predicted protein structures were downloaded from the AlphaFold Protein Structure Database (Jumper et al., 2021; Fleming et al., 2025). In cases where structures were not available or when predicting the structure of Hobbit or Bbo permutations or protein complexes, the appropriate sequence(s) was submitted to the AlphaFold Server and folded using AlphaFold3 (Abramson et al., 2024). All predicted protein structures were visualized and annotated using ChimeraX v.1.8 (Meng et al., 2023).

### Absolute quantification of mRNA transcript levels

Quantitative reverse transcription PCR and absolute quantification of transcript levels were performed as previously described (Ihry et al., 2012; Kang et al., 2017). RNA was extracted from whole animals or dissected salivary glands at the appropriate developmental stage in biological triplicate using the RNeasy Plus Mini Kit (Qiagen 74134), and cDNA was made from 400 ng of total RNA using the SuperScript III First Strand Synthesis System (Invitrogen 18080051). qPCR was performed on a Roche LightCycler 480 II with LightCycler 480 SYBR Green I Master mix (Roche 04707516001). To quantify absolute levels of target mRNAs, the cycle threshold (Ct) value for each gene in the experimental samples was compared to a standard curve of Ct values derived from the respective amplicons at known concentrations. Primer sequences were as follows: *bbo* isoform A 5’ CAG CTC TTC GAC GCC TTT GG 3’ and 5’ TGC TGT TCT GGC CTG CAG TA 3’; *bbo* isoform B 5’ CGC ACA GTC AAC ATT CAA GGC A 3’ and 5’ CGC TTC TCA CCG CTG CTT T 3’; *bbo* isoform C 5’ CGT GCG ACC ACA TTC CAC AA 3’ and 5’ CGC TTC TCA CCG CTG CTT T 3’; *bbo* isoform D 5’ AGG GTG ATT CGG GAC ATG CT 3’ and 5’ TGC TGT TCT GGC CTG CAG TA 3’. Primers were designed using the GenScript primer design tool (https://www.genscript.com/tools/real-time-pcr-taqman-primer-design-tool).

### Data availability

The data supporting the findings of this study are openly accessible in Dryad at https://doi.org/10.5061/dryad.9w0vt4bwm.

---

**Figure S1.**
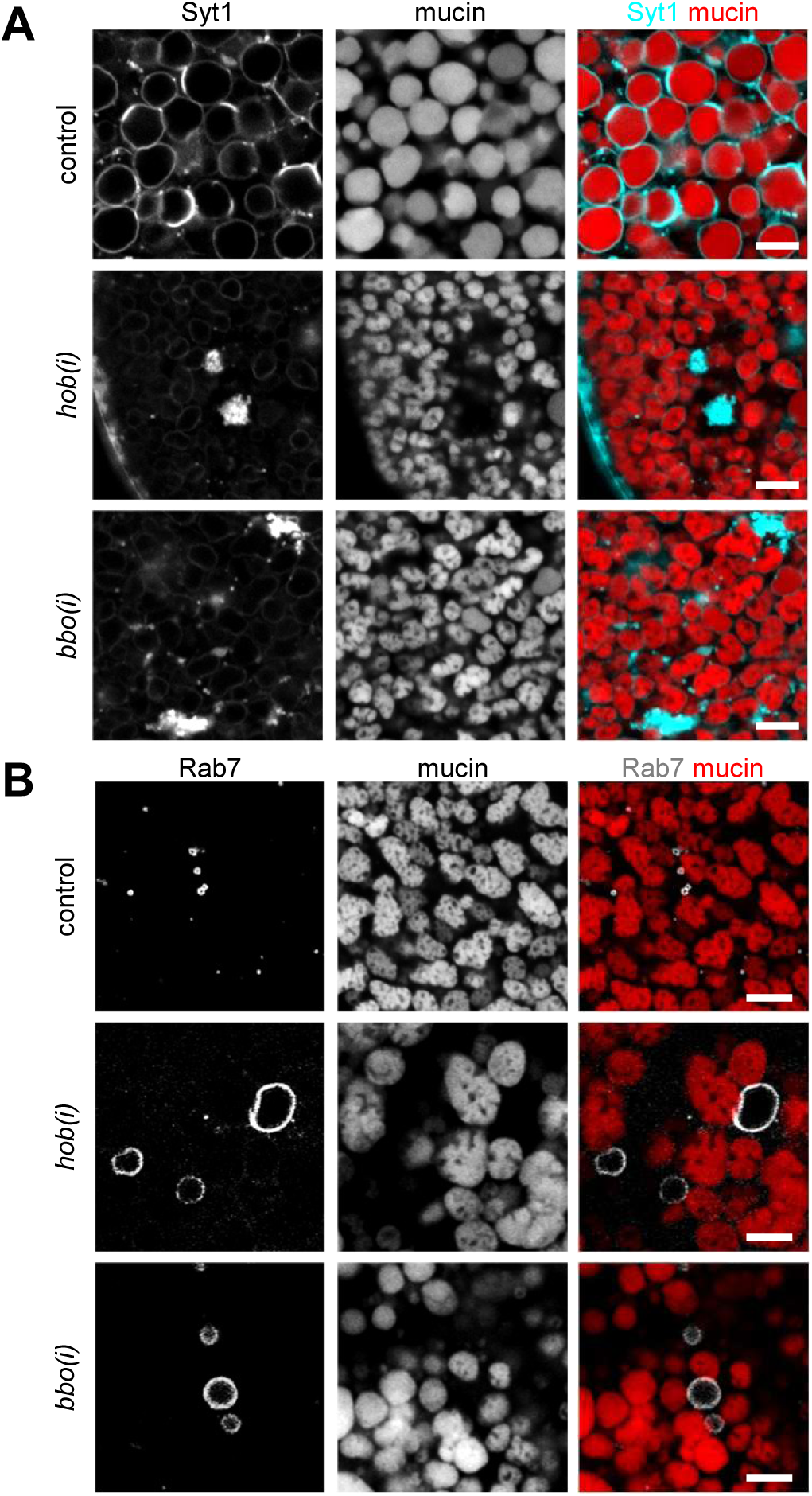
*hobbit* knockdown and *bbo* knockdown exhibit similar intracellular trafficking phenotypes. **(A)** Representative images of mucin-containing secretory granules (red) and the membrane fusion protein Synaptotagmin-1 (Syt1; cyan) in control (*Sgs3-DsRED/+; hs>/+*), *hob-RNAi* (*Sgs3-DsRED/+; hs>/UAS-hob(i)*), and *bbo-RNAi* (*Sgs3-DsRED/+); hs>/UAS-bbo(i)*) salivary glands. Syt1 is present on mucin granule membranes in control glands but is absent from granule membranes in *hob-RNAi* and *bbo-RNAi* glands. Note that Syt1 also accumulates in large clumps that are distinct from mucin granules in *hob-RNAi* and *bbo-RNAi* glands. **(B)** Representative images of mucin-containing secretory granules (red) and the late endosome marker Rab7 (gray) in control (*Sgs3-DsRED/+; Sgs3>, Rab7-EYFP/+*), *hob-RNAi* (*Sgs3-DsRED/+; Sgs3>, Rab7-EYFP/UAS-hob(i)*), and *bbo-RNAi* (*Sgs3-DsRED/+; Sgs3>, Rab7-EYFP/UAS-bbo(i*) salivary glands. Rab7-positive endosomes are small in control glands but are dramatically enlarged in *hob-RNAi* and *bbo-RNAi* glands. Note that the enlarged endosomes do not contain mucins. All images acquired from live, unfixed tissue and are representative of at least three independent experiments with *n≥*10 tissues from independent animals analyzed. Scale bars: 5 µm.

**Figure S2.**
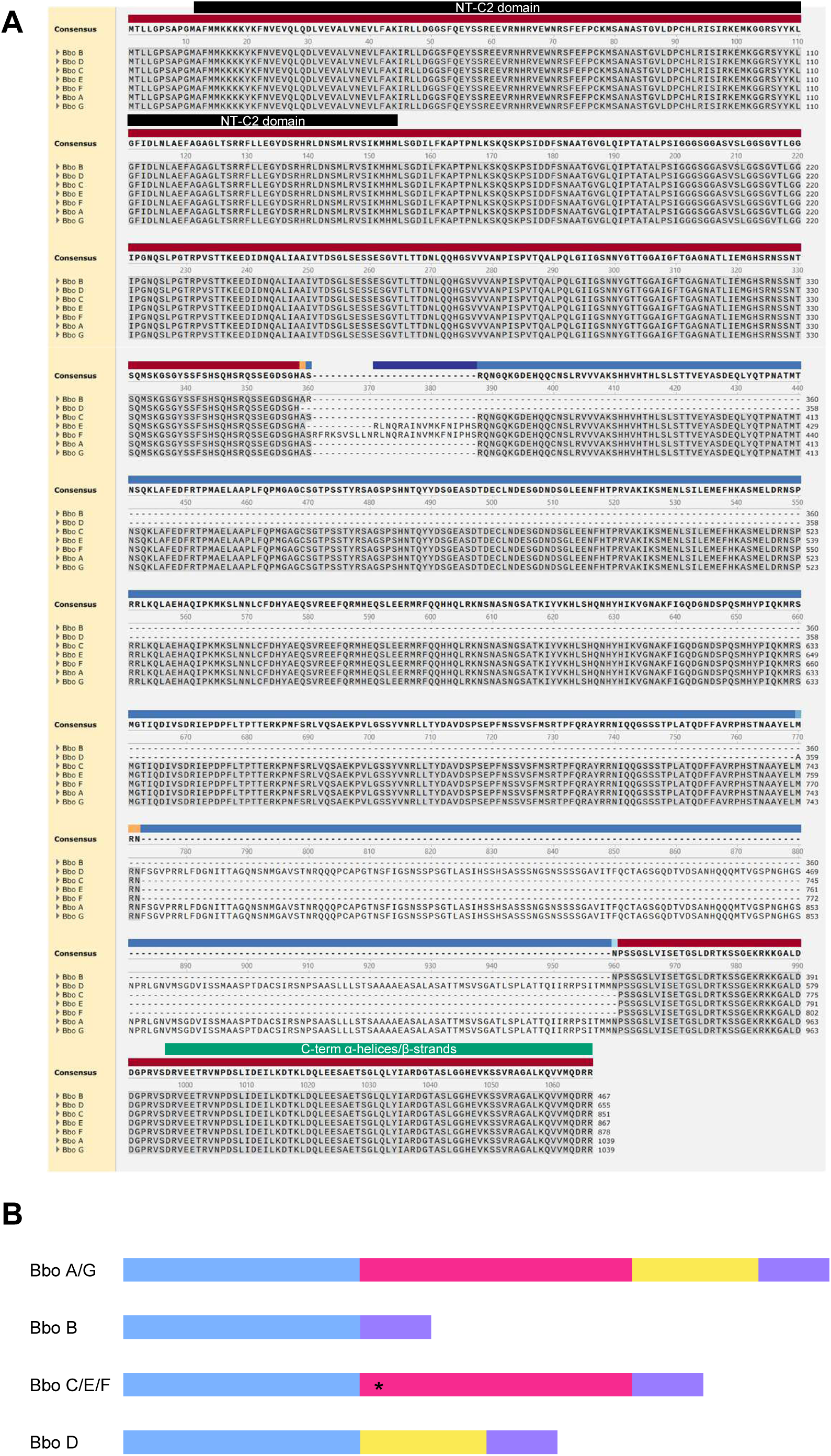
Comparison of Bbo protein isoform sequences. **(A)** Primary sequence alignment of Bbo protein isoforms; note that all isoforms contain the NT-C2 domain and C-terminal α-helices/β-strands. Alignment was generated and annotated using Snapgene. **(B)** Schematic summarizing the composition of Bbo protein isoforms based on the alignment in (A).

**Figure S3.**
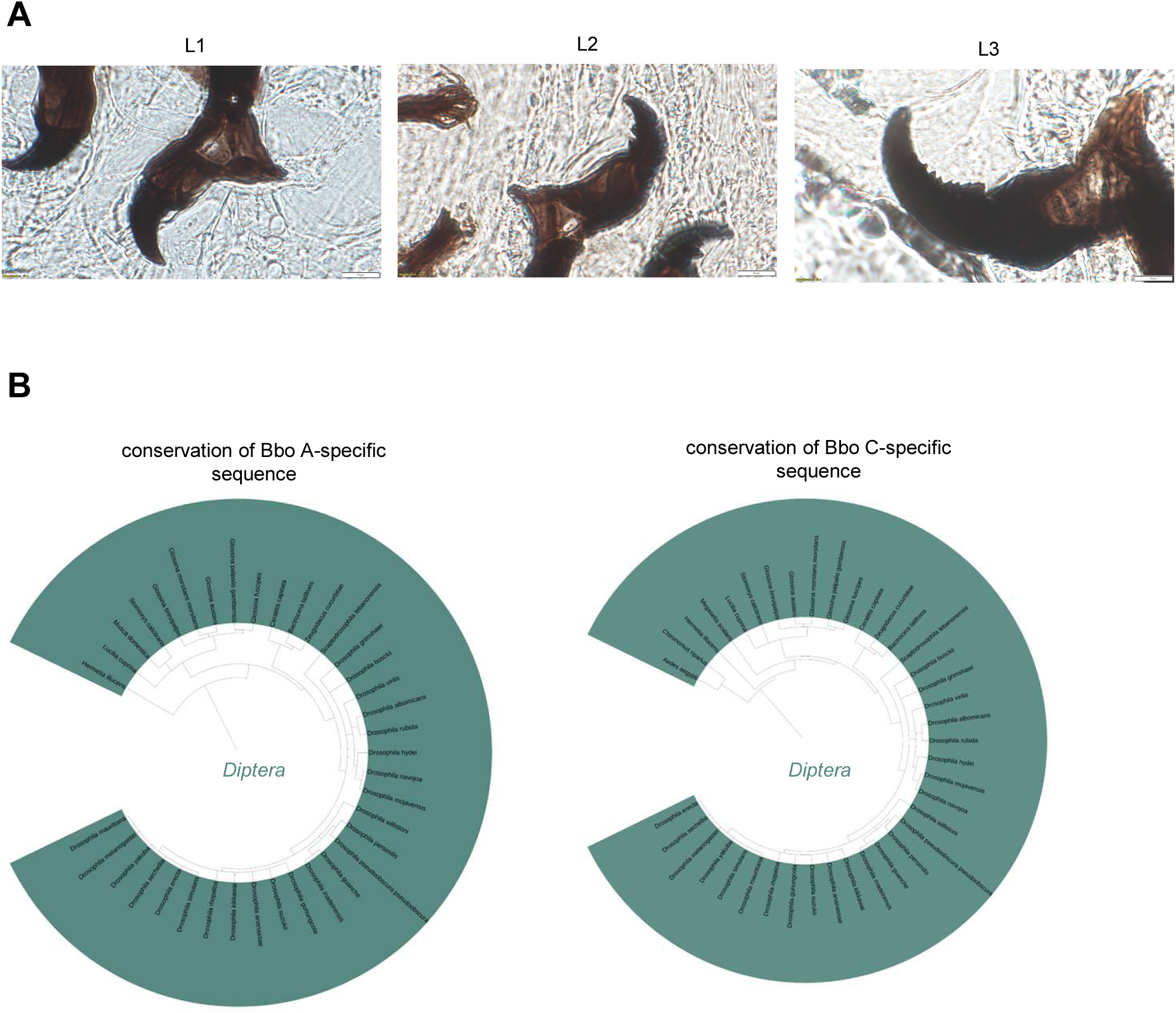
Analysis of *bbo* mutant larval phenotypes and conservation of “long” Bbo isoforms. **(A)** Representative images showing mouth hook morphology in developmentally arrested (“dead”) *bbo^KO^/Df* larvae. *n*≥10 larvae were imaged over three separate experiments. Scale bars: 10 µm. **(B)** Phylogenies showing species that contain proteins orthologous to Bbo isoform A-specific sequences (left) and Bbo isoform C-specific sequences. Note that orthologs are only found in *Diptera*; moreover, no other proteins besides Bbo orthologs were identified in this analysis. Query sequences correspond to “pink + yellow” as shown in Fig. S2B for Bbo A and “pink” for Bbo C.

**Figure S4.**
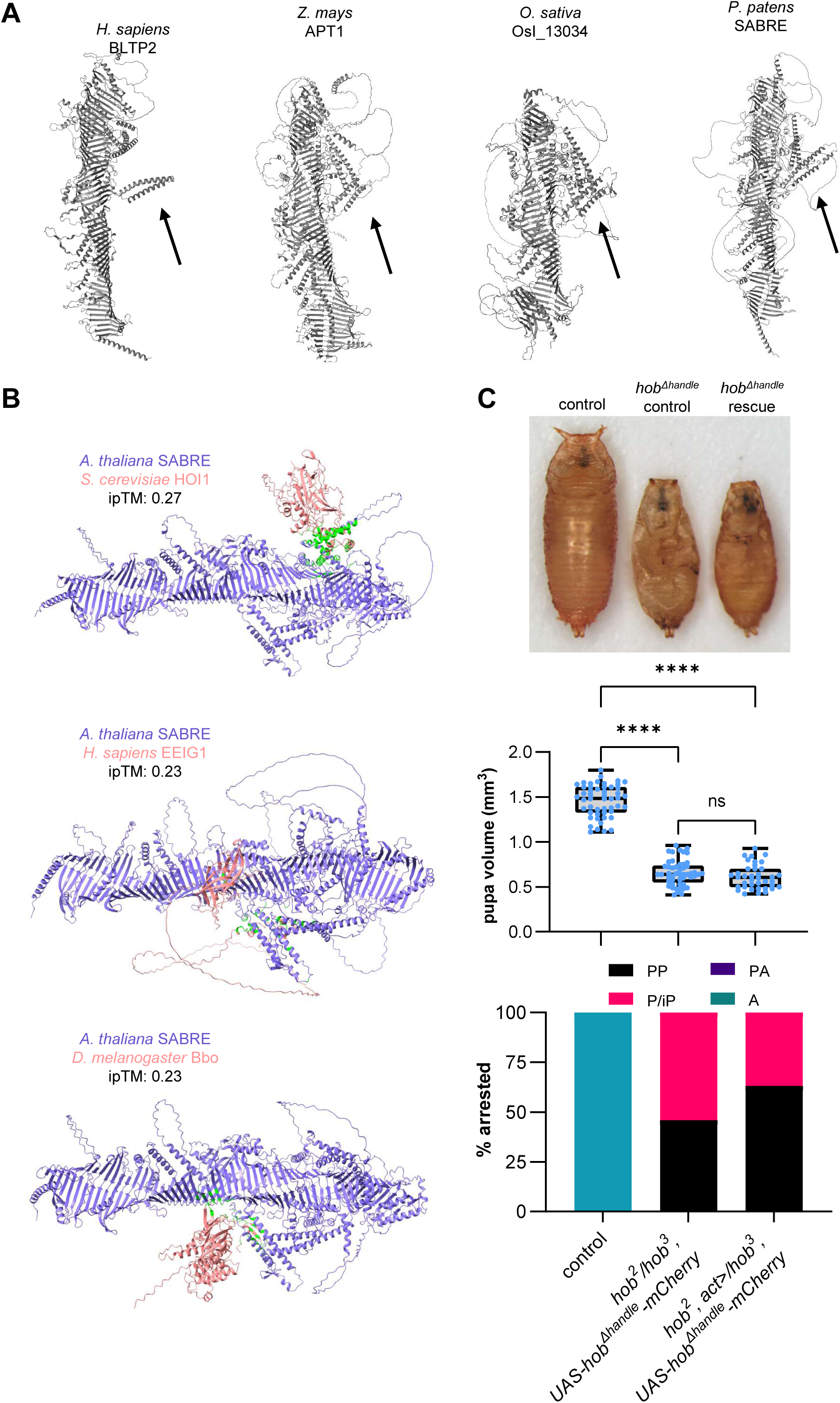
The “handle” is required for *hobbit* function. **(A)** AlphaFold predicted structures of Hobbit orthologs in *H. sapiens*, *Z. mays*, *O. sativa*, and *P. patens.* The black arrow points to the “handle” motif in each protein. **(B)** AlphaFold3 models of predicted protein-protein interactions between *A. thaliana* SABRE, purple, and *S. cerevisiae* HOI1 (top), *H. sapiens* EEIG1 (middle), or *D. melanogaster* Bbo (bottom), pink. Residues that are predicted to physically interact are colored green. **(C)** Representative image, quantification of body size (pupa volume), and lethal phase analysis in control (*w^1118^*), hob^Δhandle^ rescue control (*hob^2^/hob^3^, UAS-hob^Δhandle^-mCherry*), and hob^Δhandle^ rescue (*hob^2^, act>/hob^3^, UAS-hob^Δhandle^-mCherry*) shows that ubiquitous expression of hob^Δhandle^ does not rescue *hobbit* mutant animals; thus, the protein is not functional. Body size and lethal phase analysis were conducted on the same animals. Boxes in body size graph outline the 25th to 75th percentiles; middle line indicates the median. Whiskers extend to the minimum and maximum values. Individual data points are overlaid; statistics calculated using ordinary one-way ANOVA. *n*=50 animals for control and hob^Δhandle^ rescue control; *n*=38 animals for hob^Δhandle^ rescue. PP, prepupa; P/iP, pupa/incomplete pupa; PA, pharate adult; A, adult. **** *p*<0.0001. ns=not significant.

**Figure S5.**
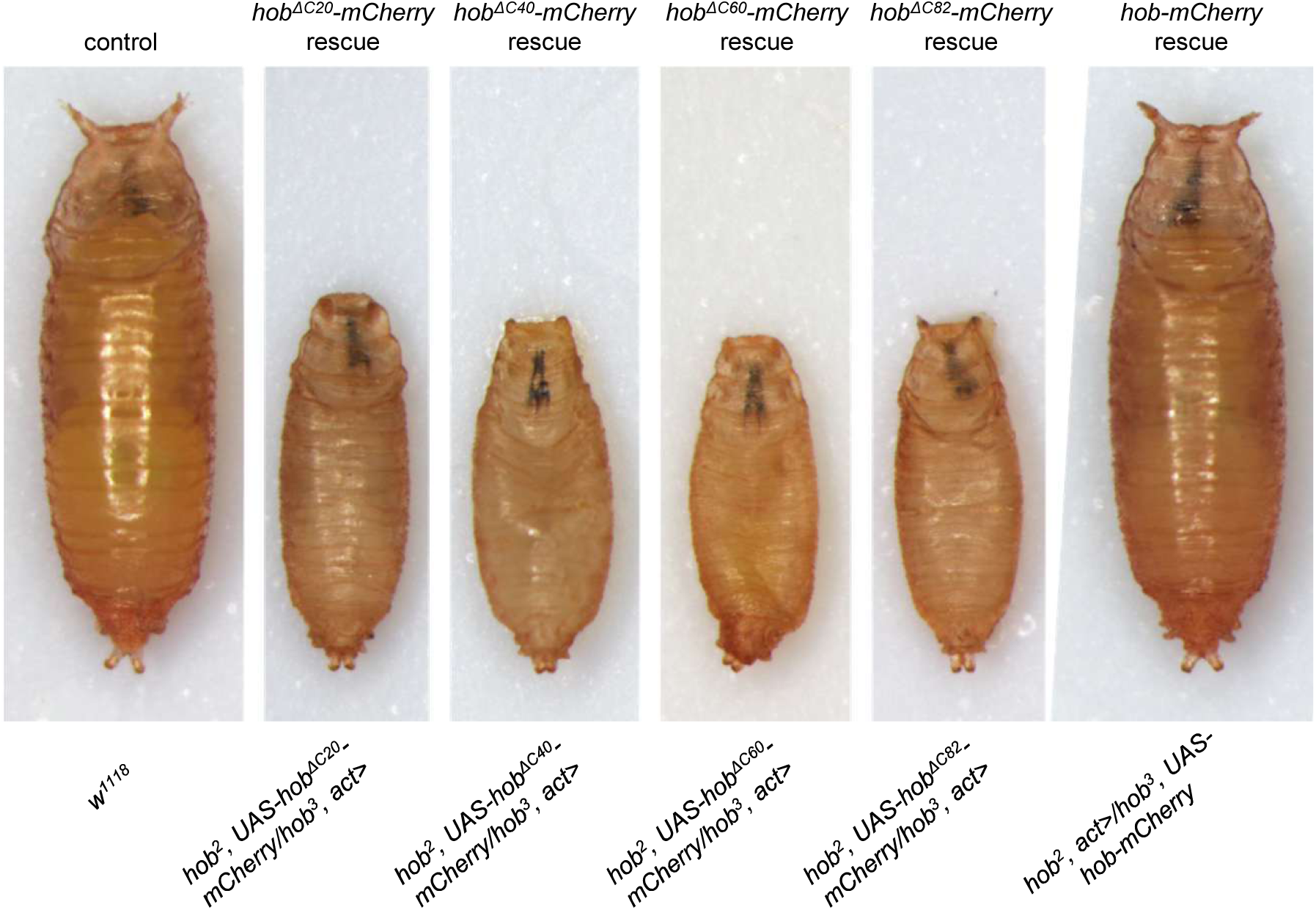
C-terminal truncations of *hobbit* do not rescue *hobbit* mutant animals. Representative images of pupa body size in control (left), rescues of *hobbit* mutant animals with ubiquitous expression of *hob^ΔC^-mCherry* truncation mutants (middle four), and rescue of *hobbit* mutant animals with ubiquitous expression of wild-type *hob-mCherry* (right). This analysis demonstrates that ubiquitous expression of wild-type Hobbit-mCherry can rescue *hobbit* mutant animals and is thus fully functional; in contrast, ubiquitous expression of the Hobbit^ΔC^-mCherry truncation mutants does not rescue *hobbit* mutant animals, indicating that C-terminal truncation disrupts *hobbit* function.

